# Personalization of hybrid brain models from neuroimaging and electrophysiology data

**DOI:** 10.1101/461350

**Authors:** R. Sanchez-Todo, R. Salvador, E. Santarnecchi, F. Wendling, G. Deco, G. Ruffini

## Abstract

Personalization is rapidly becoming standard practice in medical diagnosis and treatment. This study is part of an ambitious program towards computational personalization of neuromodulatory interventions in neuropsychiatry. We propose to model the individual human brain as a network of neural masses embedded in a realistic physical matrix capable of representing measurable electrical brain activity. We call this a *hybrid brain model (HBM)* to highlight that it encodes both biophysical and physiological characteristics of an individual brain. Although the framework is general, we provide here a pipeline for the integration of anatomical, structural and functional connectivity data obtained from magnetic resonance imaging (MRI), diffuse tensor imaging (DTI *connectome*) and electroencephalography (EEG). We personalize model parameters through a comparison of simulated cortical functional connectivity with functional connectivity profiles derived from cortically-mapped, subject-specific EEG. We show that individual information can be represented in model space through the proper adjustment of two parameters (global coupling strength and conduction velocity), and that the underlying structural information has a strong impact on the functional outcome of the model. These findings provide a proof of concept and open the door for further advances, including the model-driven design of non-invasive brain-stimulation protocols.

## 1. Introduction

The human brain is a dynamical system that can be defined as a network of networks, arranged in multiple scales and different levels of complexity. Several theoretical and computational neuroscience studies have developed whole-brain network models to explore the relationship between brain function and its underlying connectivity (Bansal et al., 2018a; Cabral et al., 2014; Deco et al., 2014). The increasing interest in finding the origin behind the structure-function relationship has led to a newly developing field known as “network neuroscience” (Bassett and Sporns, 2017), which relies on network theory to study the brain.

A growing body of evidence now suggests that brain functions emerge from interactions between specialized, spatially-segregated areas of large-scale networks. In such networks, nodes correspond to cortical or sub-cortical brain regions and edges correspond to either structural (i.e. direct connections) or functional (i.e. through synaptic interactions) couplings between these regions.

In this framework, coupled mathematical differential equations (either ordinary or partial) can be used to describe the spatiotemporal dynamics of patterns of brain activity and traveling waves (Coombes, 2005) either at the level of one node or of larger scale networks corresponding to multiple coupled nodes.

Traditionally, two main classes of models have been used to derive these differential equations.

On the one hand, spiking neuron models such as the Hodgkin-Huxley model (Hodgkin and Huxley, 1990) describe the dynamics of individual neurons. On the other, neural mass models (NMM) such as the Wilson-Cowan model (Wilson and Cowan, 1972) provide effective theories of neural systems. The former, more detailed class of models is appropriate for representing single cell recordings in animals or brain slices, but their state variables do not directly capture the functional activity recorded with macroscopic level techniques such as Electroencephalography (EEG), Magnetoencephalography (MEG) or mesoscopic Local Field Potential (LFP) measurements. In contrast, NMMs are more relevant when modeling brain activity at larger spatial and temporal scales since they describe the mean activity of whole neural populations. Despite providing a lower level of detail, they are able to represent the physiology of the brain: their parameters emerge from microscopically measurable quantities, such as dendritic time constants and mean excitatory/inhibitory post-synaptic potentials.

Depending on the data available or the complexity that we want to model, network nodes can represent either single cells, cortical patches, or whole brain areas (see an extended review by Breakspear (2017) for a detailed discussion on the choices of dynamical equations). Accordingly, network edges are to describe appropriately the links between nodes. E.g., coupling strength is often defined in proportion to the number of white matter tracts (structural connectivity) between brain areas to model whole-brain dynamics (Honey et al., 2007, 2009; T. Proix et al., 2017; Deco et al., 2018), using the well-known human connectome (Hagmann et al., 2007). However, functional connectivity has also been used to define these links (Bassett DS, 2009; Feldt et al., 2011; Bassett and Sporns, 2017).

Bansal et al. (2018b) and Aerts et al. (2016) review recent research on personalized whole-brain models. The former is related to the study of structure-function relationship in human brains and the second focuses on the impact of network lesions. The majority of the studies cited in these reviews only use structural connectivity brain data derived from DTI to personalize whole-brain models. Moreover, most of the models in those studies are based on static network structures with fixed connection strength and time delays, failing to reproduce some meaningful features on brain dynamics that may vary between subjects.

A few recent studies have focused on personalizing dynamical model parameters based on subject specific data. For example, Finger et al. (2016) and Cabral et al. (2014) optimized the coupling gain and the mean conduction velocity of a network of Kuramoto oscillators based on the correlation of functional connectivity (FC) profiles between simulated and real activity. A common feature of this type of resting-state computational models is that slow signal fluctuations (*<* 0.1Hz) appear when the system operates at the edge of bifurcations, due to specific combination of model parameters (Honey et al., 2007, 2009; Deco et al., 2009; Cabral et al., 2011, 2014). When operating close to bifurcations, the dynamics of the network reveal patterns that are shaped by the underlying anatomical connectivity. Nevertheless, the relationship between structural and functional connectivity derived from EEG in the resting-state has not been deeply addressed in computational models.

Although electrophysiology data has been used to reveal ongoing activity of the whole-brain since the beginning of the past century (Haas, 2003), the first description of resting-state networks was provided by functional MRI (fMRI)/Position Emission Tomography (PET) recording techniques, in studies that revealed correlated slow fluctuations (*<* 0.1 Hz) across regions (Biswal et al., 1995; Raichle et al., 2001). Likewise, the envelope of the amplitudes of alpha (8-14 Hz) and beta (14-30 Hz) frequency oscillations of the electrophysiological signals exhibit, when observed at slow time scales (*<* 0.1 Hz), correlation patterns similar to those of fMRI signals (Hipp et al., 2012; Brookes et al., 2011). Furthermore, there is evidence that there is a strong dependence of slow fluctuations of resting-state networks (*<* 0.1Hz) on white matter pathways — indicating that the anatomical connectivity could shape the functional connections among brain areas (Honey et al., 2009; Hagmann et al., 2008).

In this study we propose to represent the human brain as a network of biologically realistic neural mass models, and then personalize such a virtual brain with subject-specific data (Figure 1), both biophysical (MRI) and physiological (EEG) characteristics at the individual level, and, if available, also structural information of the subject (from DTI). We call this personalized *hybrid brain model (HBM)* to highlight that it can encode both biophysical and physiological characteristics of the individual human brain.

**Figure 1:**
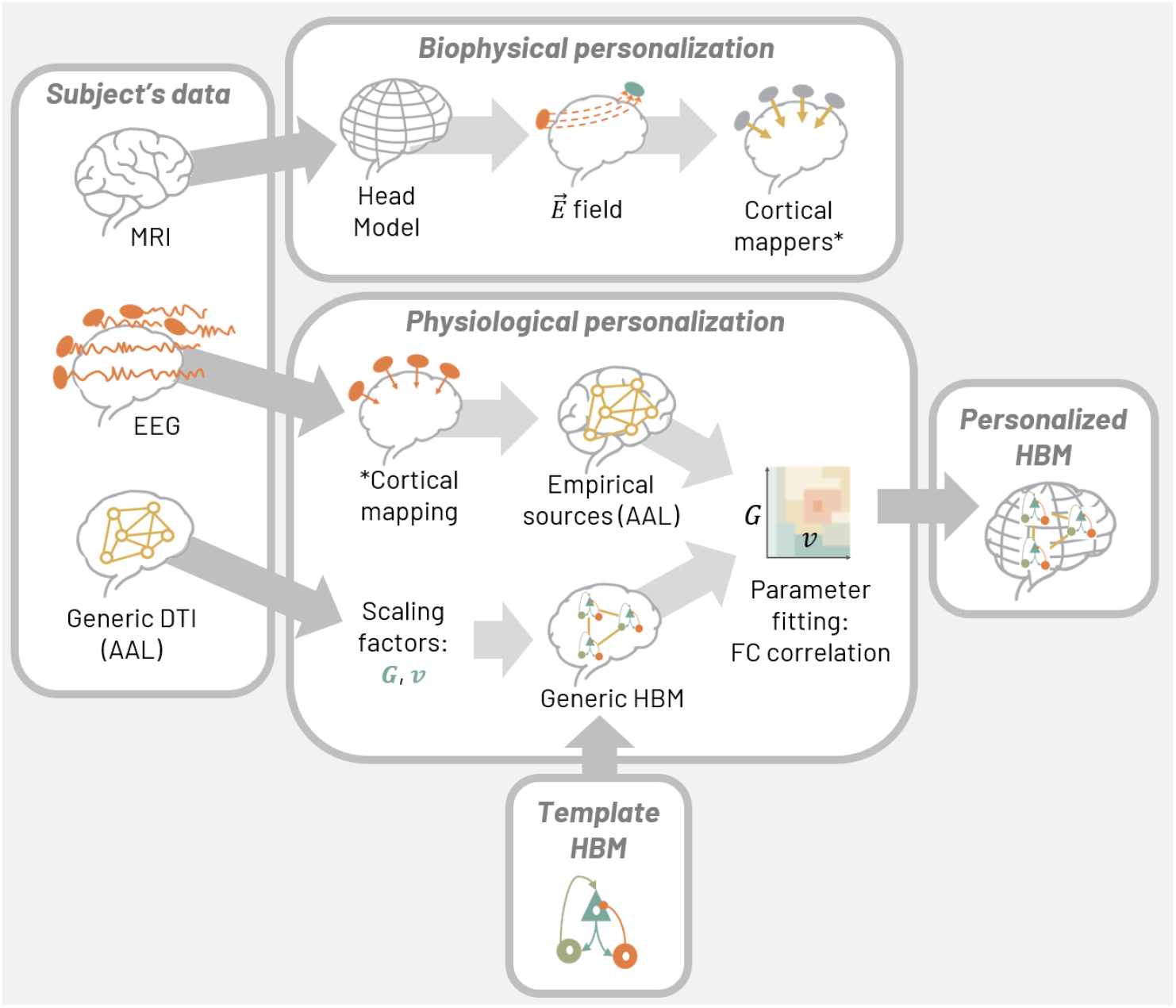
Personalized Hybrid Brain Model generation using subject MRI, EEG and DTI data. <u>Biophysical personalization</u>: a realistic head model of electrical conduction is generated using MRI data and finite element modeling, solving Poisson’s equation. The realistic head model is used for cortical mapping (*) of the subject’s EEG into cortical mesh dipole activity (Ruffini, 2015), and may also be used for tCS modeling and optimization. <u>Physiological personalization</u>: a template HBM is created and registered to the subject cortex (realistic head model). Cortically mapped EEG data is then averaged over selected AAL areas and used to create a personal functional connectivity (FC) profile. <u>Personalized HBM:</u> the personal FC profile is compared to the one produced by the HBM adjusted using a family of parameters. The result of the comparison of data- and model-derived FC quantified by cross-correlation is used to adjust HBM parameters for best fit.

We will first describe our construction of a *template* HBM, composed by a network of realistically interconnected Jansen and Rit (Jansen et al., 1993) NMMs which can simulate mesoscopic and macroscopic activity of the brain. We have also chosen this particular NMM because it can be tuned to simulate healthy or physio-pathological electric signals (Wendling et al., 2016; Kurbatova et al., 2016) and its possible response to interventions like transcranial current/magnetic stimulation (tCS, TMS) (Ruffini et al., 2013; Merlet et al., 2013; Molaee-Ardekani et al., 2013; Kunze et al., 2016; Modolo et al., 2018) or drugs (Liang et al., 2015; Kurbatova et al., 2016). The main reason behind developing mesoscopic models of neural dynamics at the scale of whole brain is that experimental data is recorded at this scale (EEG, MRI). Therefore, network edges in the proposed HBM represent anatomical links (structural connectivity) between brain areas. Since we do not account here for subject-specific DTI, we use an averaged and scaled DTI data, as in in Cabral et al. (2014). Inspired by the work of Molaee-Ardekani et al. (2013) and Merlet et al. (2013), we fit the key parameters of the template HBM in two different perspectives: (1) biophysically, using subject-specific MRI to create human head models (Miranda et al., 2013, 2018) and (2) physiologically, correlating the simulated FC with the one coming from cortical mapped electrophysiology data (EEG) (Figure 1).

Biophysical aspects of the HBM are essential to couple physical phenomena with physiology. Realistic, finite element models of the head can represent, for example, the electric fields and scalp potentials generated by the electrical activity of the brain, or the alterations in membrane potential generated by non-invasive brain stimulation (Abea et al., 2018). Thus, they can be used to project the neural dynamics of coupled NMMs into measurable EEG, and to represent the effects of transcranial current/magnetic stimulation (tCS/TMS) on neural dynamics.

The physiological personalization process (i.e., parameter fitting) is based on the comparison of the measured functional connectivity (FC) in source space (cortical dipole activity from cortically mapped EEG, Ruffini (2015)) and the one derived here from the HBM (see Figure 1). Mathematically, this comparison is quantified by the cross-correlation coefficient between the model FC and the empirical one. The source space distribution is averaged over the same parcellation atlas used to define the nodes in the HBM. The FC estimate is based on the EEG filtering pipeline of Cabral et al. (2014), which takes into account the fact that resting state blood-oxygen-level dependent (BOLD) signal fluctuations are driven by slow modulations in the power (and therefore, amplitude) of brain oscillations in different frequency ranges.

In this work, using this definition of a HBM, we first explore the relationship between the underlying structural connectivity and the functional outcome of the HBM. Although our aim is to provide a proof of concept of the method, we also study how well our modeling framework can fit subject-specific data based on the correlation between measured and simulated FC profiles, explore how tightly model parameters are determined by the data, and comment on the role of inter-subject variability. Finally, we discuss potential uses of HBMs in the clinical domain and future research directions.

## 2. Materials and Methods

### 2.1. Participant information and data

In this study we analyzed EEG and MRI from two right-handed healthy male volunteers (aged 22 and 24 years old), collected as part of a study about the identification of novel biomarkers for Neuropsychiatric disorders based on individual patterns of response to external perturbation (“Perturbation-based Biomarkers”) supported by the Harvard-MIT Broad Institute. We have labeled them as subject 55 and subject 60. Experimental protocols and voluntary participation procedures were explained to all participants before they gave their written consent. Experimental protocols herein conformed to the Declaration of Helsinki, and the Institutional Review Board of the Beth Israel Deaconess Medical Center (BIDMC, Boston, MA, USA) approved the study protocols. The study included concurrent Transcranial Magnetic Stimulation (TMS) and EEG recording, looking at individual patterns of EEG propagation in response to single TMS pulses delivered over cortical regions defined using individual fMRI/MRI data. Details on the EEG and MRI data are provided below.

A T1-weighted anatomical MRI scan was acquired on a 3T MRI system (GE Healthcare, Ltd., United Kingdom) using a 3D spoiled gradient echo sequence: 162 axial-oriented slices for whole-brain coverage; 240-mm isotropic field-of-view; 0.937-mm *×* 0.937-mm *×* 1-mm native resolution; flip angle = 15^*o*^; TE/TR *≥* 2.9/6.9 ms; duration *≥* 432 s.

EEG was recorded while participants were comfortably seated in an adjustable chair, with their eyes open (EO) while fixating a cross-air on a PC monitor, and then with the eyes closed (EC). Resting-state EEG was recorded for three minutes in each condition (EO, EC). Whole scalp 64-channel EEG data was collected with a TMS compatible amplifier system (actiCHamp system, Brain Products GmbH, Munich, Germany) and with electrodes placed in accordance with the extended 10–20 international system. EEG data were online referenced to the Fp1 electrode. Electrode impedances were maintained below 5 kΩ at a sampling rate of 1000 S/s. EEG signals were digitized using a BrainCHamp DC amplifier and linked to BrainVision Recorder software (version 1.21) for online monitoring. Digitized EEG electrode locations on the scalp were also co-registered to individual MRI scans using Brainsight^TM^ TMS Frameless Navigation system.

In this paper we did not use individualized tractography data to define the connectivity and delay matrices. Instead, for simplicity, we use a structural connectivity matrix derived from Diffusion Tensor Imaging (DTI) of 21 healthy participants, averaged among 78 cortical areas in the AAL atlas (see Table B.1), as described in (Cabral et al., 2014), to define a prototypical coupling matrix, *C^ij^*. The delay matrix across nodes 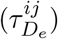 is estimated from the euclidean distance, *D_e_*, between the center of gravity of the parcellated areas (see next section for a detailed description of the model parameters *C^ij^*, and 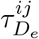).

### 2.2. Template hybrid brain model (HBM)

The template HBM is defined here as a networked neural mass model, with biophysical aspects, node positions, connectivities and conduction delays unspecified. As discussed above, we work here with an extension of Jansen and Rit model, that encodes network aspects in new parameters, the connectivity and delay graph adjacency matrices. See Appendix A for a detailed mathematical formulation of Jansen and Rit NMM. Jansen and Rit (Jansen and Rit, 1995) developed a NMM of cortical columns based on the thalamus model of Lopes da Silva et al. (1974) and on the post-synaptic-potential (PSP) transformation function that van Rotterdam et al. (1982) designed. This model simulates a population of pyramidal cells located in layer V of the cortex that receive inhibitory inputs from a sub-population of interneurons, dendrite targeting cells. They also receive excitatory feedback either from a population of excitatory interneurons or other pyramidal cells located in the same cortical column.

The Jansen and Rit NMM of cortical columns has three main state variables, the average membrane potential of each of the subpopulations, *y*_0_ for the pyramidal cells and *y*_1_, *y*_2_ for the excitatory and inhibitory interneurons, respectively. The main output of the model is the average membrane potential of the pyramidal cell population, as the sum of the potential of these cells is thought to be the source of the potential recorded in the EEG.

In order to generate a network of such neural populations, we assume here for simplicity that all the connections across columns are from pyramidal to pyramidal populations and thus they are always excitatory. The average firing rate of action potentials from the pyramidal cells of one population, *x*(*t*), is used as an excitatory input to the rest of populations in the network. However, as neural populations represent distinct and distant areas of the cortex, new parameters need to be introduced to account for the structural connection strength among the areas and for the transmission delays associated with each connection. We also assume here that the same cortical column model can be used for all the different cortical areas of the brain, since the basic neuronal architecture of the cortex is similar throughout its areas.

Following the approach in Wendling and Chauvel (2008), a connectivity matrix *K^ij^* is used to define the degree of coupling between population *i* and *j*. In this study, *K^ij^* = *G • C^ij^*, where *C^ij^* is the normalized and symmetric (divided element wise by its maximum value) structural matrix derived from averaged DTI (discussed in section 2.1), and *G* is a scaling factor to be estimated by fitting the model with subject-specific data.

The other factor not in the original Jansen and Rit NMM and that we will use for model fitting is the transmission velocity *v* of signals across populations. The associated delay matrix will be taken as a function of the Euclidean distance matrix, 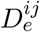, between the center of mass of the aforementioned AAL areas, and the conduction velocity *v*, 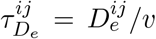 For simplicity, this conduction velocity is assumed to be homogeneous across the brain (Deco et al., 2009).

The coupled Jansen and Rit model that we present in this study therefore includes a new input to the pyramidal cells of each population model that reflects the connectivity across areas, *K^ij^*, and which is proportional to the density of white matter tracts between areas, and with a transmission delay, *τ_D_*, that is proportional to the euclidean distance among them and inversely proportional to conduction velocity *v*. The equations for each population *i* are then given by

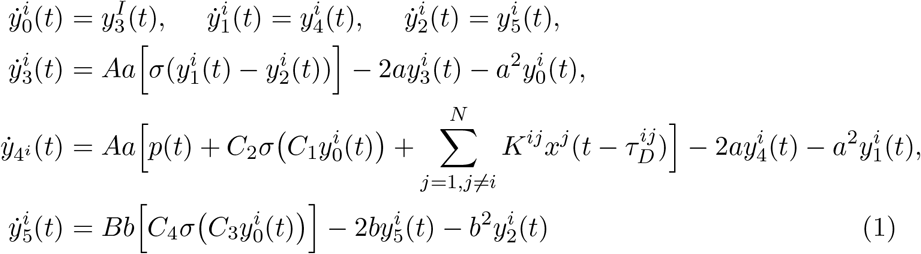

with the column input from another population *x^j^* included in the equation for 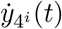 (Figure 2).

**Figure 2:**
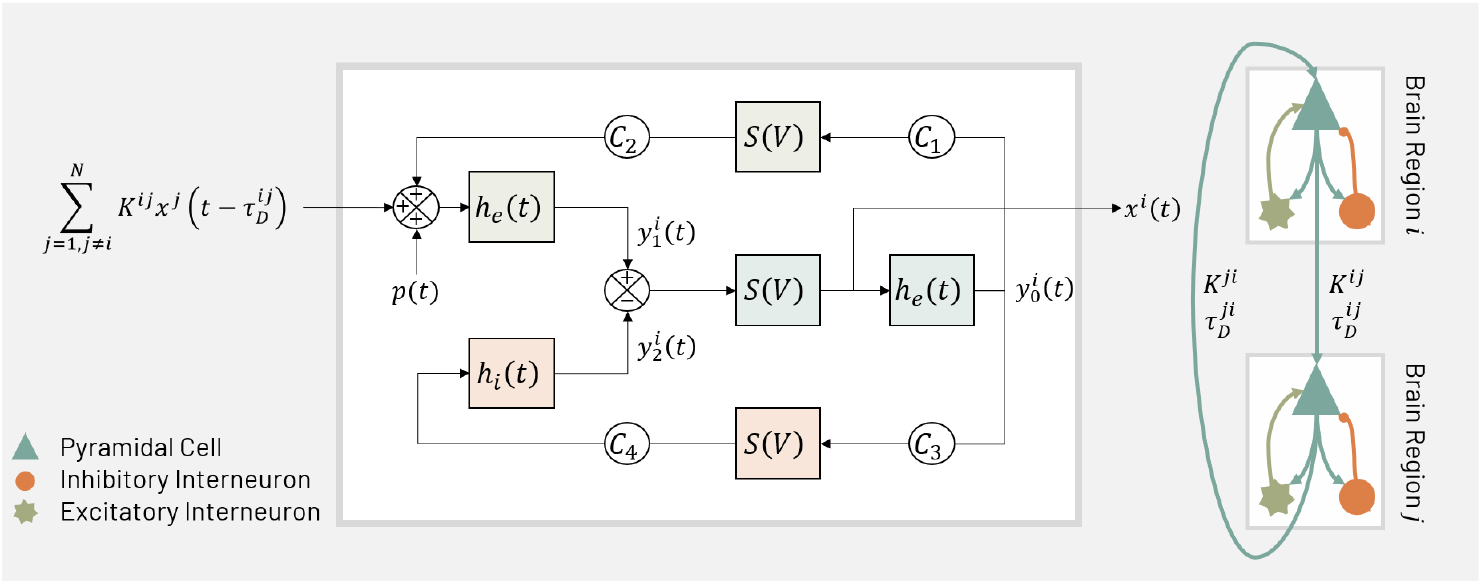
Physiological model: extension of Jansen and Rit NMM. The original Jansen and Rit model (Jansen et al., 1993) (white box, see description in Appendix A) is extended to reflect connectivity among NMMs using the matrix *K^ij^*, which represents the number of white matter tracts (average structural connectivity matrix), and to take into account the transmission delay among areas, with 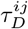 the delay taken here as proportional to the Euclidean distance.

### 2.3. From template to personalized HBM: biophysics

We next discuss how the generic HBM template is personalized using individual neuroimaging data. A summary of the pipeline followed is illustrated in Figure 3.

**Figure 3:**
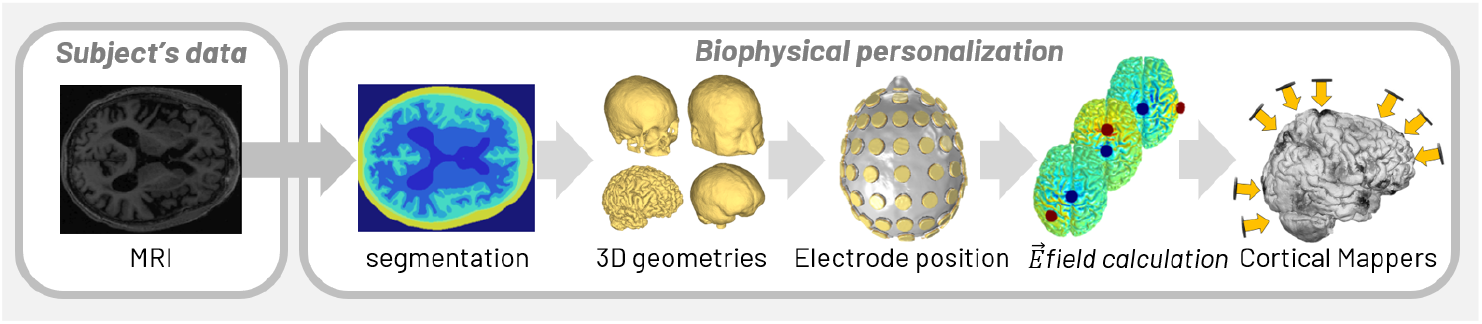
Biophysical personalization and cortical mapper generation. From the subject-specific T1 weighed images, we segmented and generated a volume mesh for each of the tissues: scalp, skull, CSF, GM and WM. Then we placed the electrodes and computed the electric field distribution over the surface mesh for each pair of electrodes. This information is then used to create the personalized cortical mappers using reciprocity principles, as described in Ruffini (2015).

#### 2.3.1. Realistic head model and calculation of lead fields

The realistic head models were built from the T1-weighted images of the same subjects that underwent the EEG recordings. The images were segmented into scalp, skull, air cavities, cerebrospinal fluid (CSF), grey-matter (GM) and white-matter (WM). Segmentation of the scalp, skull and CSF (including ventricles) was performed in Matlab (v2018a, https://ch.mathworks.com/products/matlab.html) using SPM8 with the MARS toolbox. Segmentation of the GM and WM tissues was performed in Freesurfer. The resulting segmentation masks were combined in Matlab using custom scripts. Triangulated surface meshes were then produced of all the tissues using the Iso2Mesh toolbox (Fang and Boas, 2009). Some changes in the surfaces meshes were conducted to prevent tissue intersections and further smooth the resulting surfaces. Models of the electrodes used in the EEG recording were then placed on the scalp surface according to the positions defined in the 10/10 EEG system. Those positions were defined based on manually placed anatomical landmarks (*inium, nasium* and pre-auricular left and right points). A finite element volume mesh of the final head model with the electrodes was created using Iso2Mesh. The mesh comprises around 3.5 million tetrahedral elements.

The models were used to calculate the E-field distribution induced in the brain with all bipolar montages having one of the electrodes as the anode (+1 mA) and Cz as a common cathode (−1 mA). It is possible to show that the E-field in any given montage involving these electrodes can be calculated as a linear superposition of the E-field in these files (scaled by the currents in the electrodes), Ruffini et al. (2014). Calculation of the electric field (*E*-field or lead field) was performed in Comsol (v5.3a, www.comsol.com) using second order Lagrange elements and an iterative algorithm (Conjugate Gradients solver, with an Algebraic Multigrid pre-conditioner) to solve Laplace’s equation,

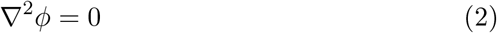

with appropriate boundary conditions Miranda et al. (2013). Once this equation is solved, the E-field can be obtained by taking the gradient of the electrostatic potential: 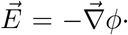 The head model volume mesh was then imported into Comsol, where tissue electrical conductivities were assigned: 0.33 S/m for the scalp, 0.008 S/m for the skull, 1.79 S/m for the CSF, 0.4 S/m for the GM, 0.15 S/m for the WM and 10^*−*5^ S/m for air, Miranda et al. (2013). The electrodes were represented as conductive gel, 4.0 S/m. After the calculation, we exported the component of the electric field normal (orthogonal) to the cortical surface 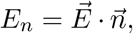 where 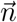 is a vector normal to the cortical surface and pointing inwards). Finally, we down-sampled the grey matter mesh and electric field distribution over the surface to a 25% of its original size (~ 40,000 mesh points) to save computational time in what follows.

#### 2.3.2. Personalized cortical mapper

EEG signals are thought to arise from the sum of post synaptic potentials occurring on pyramidal cells, which are located in the grey matter and oriented perpendicular to its surface (Nunez and Srinivasan, 2006). Accordingly, to construct a cortical mapper from electrode to source space, we assumed that dipole generators of EEG are located on the cortical surface and oriented orthogonally to it. In order to find the dipole distribution over the grey matter surface, we computed and used a personalized cortical mapper based on the reciprocity theorem (derived by H.Helmholtz in 1853, see Plonsey (1969)) and used to invert EEG as described in Ruffini (2015, 2016). Briefly, starting from a realistic forward model of electric currents generated by transcranial stimulation, a forward model of EEG and the associated cortically-bound inverse model are derived.

### 2.4. From template to personalized HBM: physiology

#### 2.4.1. From EEG to cortical source space

The preprocessing of EEG data was performed offline using a combination of the EEGLab toolbox (Delorme and Makeig, 2004) and custom scripts in Matlab 2016a (Mathworks, USA). First, data were filtered for power line noise using a notch filter (55-65 Hz) and additional low-(55 Hz) and high-pass (1 Hz) filters were applied using a zero-phase second-order Butterworth filter. Subsequently, we rejected manually excessively noisy epochs by visual inspection, and interpolated semi-automatically detected bad channels. The remaining data was re-referenced to Cz, so we could perform the personalized cortical mapping (see sections below). After re-referencing, we selected 1 min of recordings in order to compare it with the simulated data. The cortical mapper derived from the biophysical models (Section 2.3.2) was then used to to project the EEG data from electrode to cortical source space.

#### 2.4.2. Parameter fitting from functional connectivity

After source inversion of EEG data into the subject-specific grey matter mesh, we averaged the activity of each of the nodes within each of the 78 cortical areas of the AAL atlas (Tzourio-Mazoyer et al., 2002), which can be compared with the simulated data of the model (a list of the cortical AAL areas and the ordering among the matrices is provided in Appendix B). The AAL template was used to parcellate the brain originally into 116 regions, but we restricted the parcellation to 78 cortical areas according to Achard (2006) and Gong et al. (2009). We performed such reduction because we use subject specific 3D grey matter meshes and when performing the automatic parcellation into the 116 AAL areas, some of the subcortical areas are not represented in any of the mesh points. In the end, the model includes 78 cortical areas, 39 for each of the hemispheres (see Table B.1).

In order to partially eliminate zero-lag correlations introduced by cortical mapping (due to the ill-posed nature of the inverse problem), we removed the common mode component of the time series in each area.

Following the pipeline established in (Cabral et al., 2012), and considering that BOLD signal fluctuations observed in the brain at rest are associated to slow fluctuations in the power of neuronal oscillations occurring in a particular frequency range (Hipp et al., 2012; Brookes et al., 2011), we band-pass filtered (BPF) the real data and the HBM output (time courses of each of the 78 areas) into 5 different frequencies: delta (*<* 4Hz), theta (4-8 Hz), alpha (8-14 Hz), beta (14-30 Hz), slow gamma (30-58 Hz) (following the Recommendations for the Practice of Clinical Neurophysiology: Guidelines of the International Federation of Clinical Physiology (EEG Suppl. 52)).

Since the low-frequency component of the envelope fluctuations has been found to be optimal for measuring spontaneous functional connectivity (Hipp et al., 2012; Brookes et al., 2011), we first extracted the amplitude envelope using Hilbert transform and then low-pass filtered (LPF) the time courses with a cut-off frequency of 0.5 Hz.

Finally, the Pearson Correlation Coefficient (PCC) between areas was extracted, representing the final FC profile of the HBM output or of real data in the form of a network adjacency matrix.

The last step in personalization is to fix the two free parameters in our HBM model: conduction velocity *v* and global coupling strength *G*. In order to do this we searched for the parameter combination that led to maximal correlation of the real and simulated FC adjacency matrices. That is, a FC-correlation error cost function was used to optimize our two parameters by comparing the PCC for each pair of parameters *G* and *v*.

Figure 4 provides a diagram of the pipeline to used extract the FC profile for each combination of model parameters.

**Figure 4:**
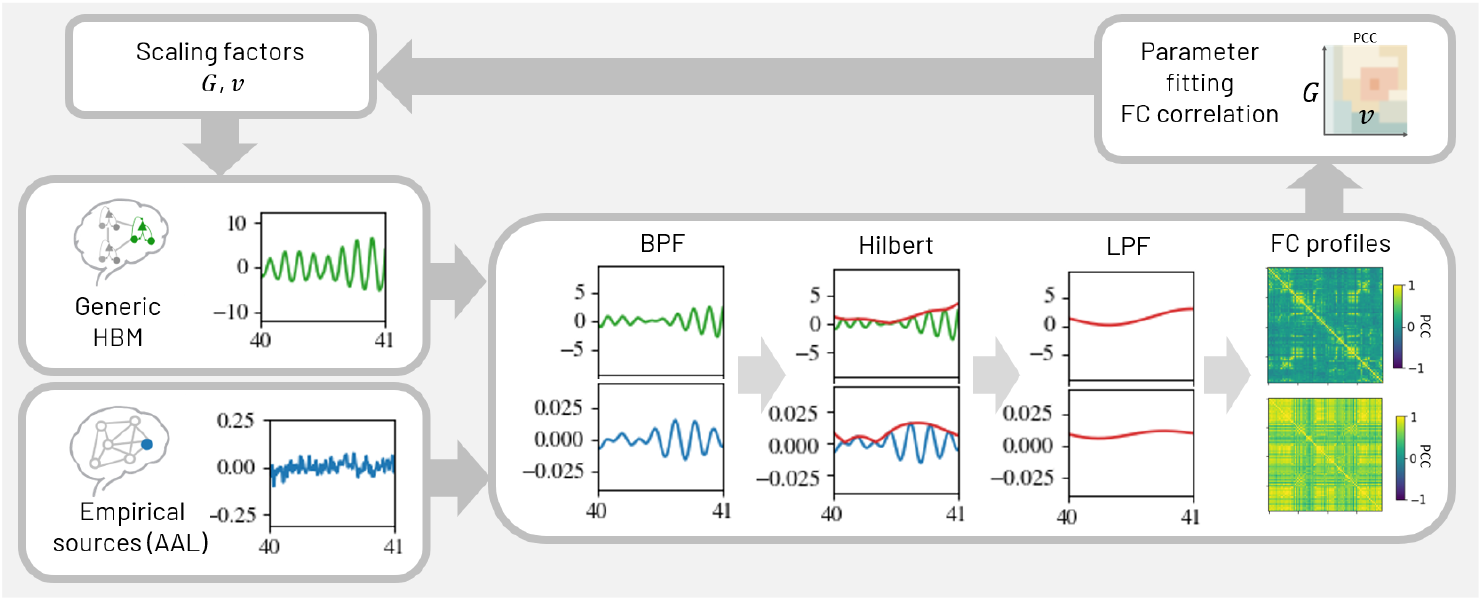
Parameter fitting based on the correlation coefficient between real and simulated FC matrix for each combination of model parameters. The FC profiles were generated by band pass filtering the simulated (green) and real (blue) data, computing the Hilbert transform (envelope) of the signal and finally low pass filtering it (*<*1 Hz). We computed the correlation of this signal across cortical areas to extract real and simulated FC matrices. Finally, we calculated the correlation between real and simulated FC matrices and identified which of the parameters produced the best fit.

## 3. Results and Discussion

### 3.1. Correlation with the underlying connectivity matrix

The relationship between functional and structural connectivity is not trivial and has been the subject of investigation by several theoretical and computational research groups over the last decade (Deco et al., 2009; Honey et al., 2007; Ponce-Alvarez et al., 2015). These efforts have shown, using whole-brain network models, how anatomic structure can shape spontaneous brain activity on slow fluctuations, displaying consistent patterns of functional connectivity.

In Figure 5 we provide correlation maps derived from the comparison of the generic HBM’s underlying structural connectivity matrix (which arises from average DTI data and is used to connect the NMMs in our HBM) with the FC profile of the HBM for different combination of model parameters: coupling gain, *G*, and mean velocity, *v*. As Figure 5 shows, the underlying connectivity information of our model has indeed a strong influence on the FC profile, achieving maximum values of *PCC* = 0.74 for specific parameters.

**Figure 5:**
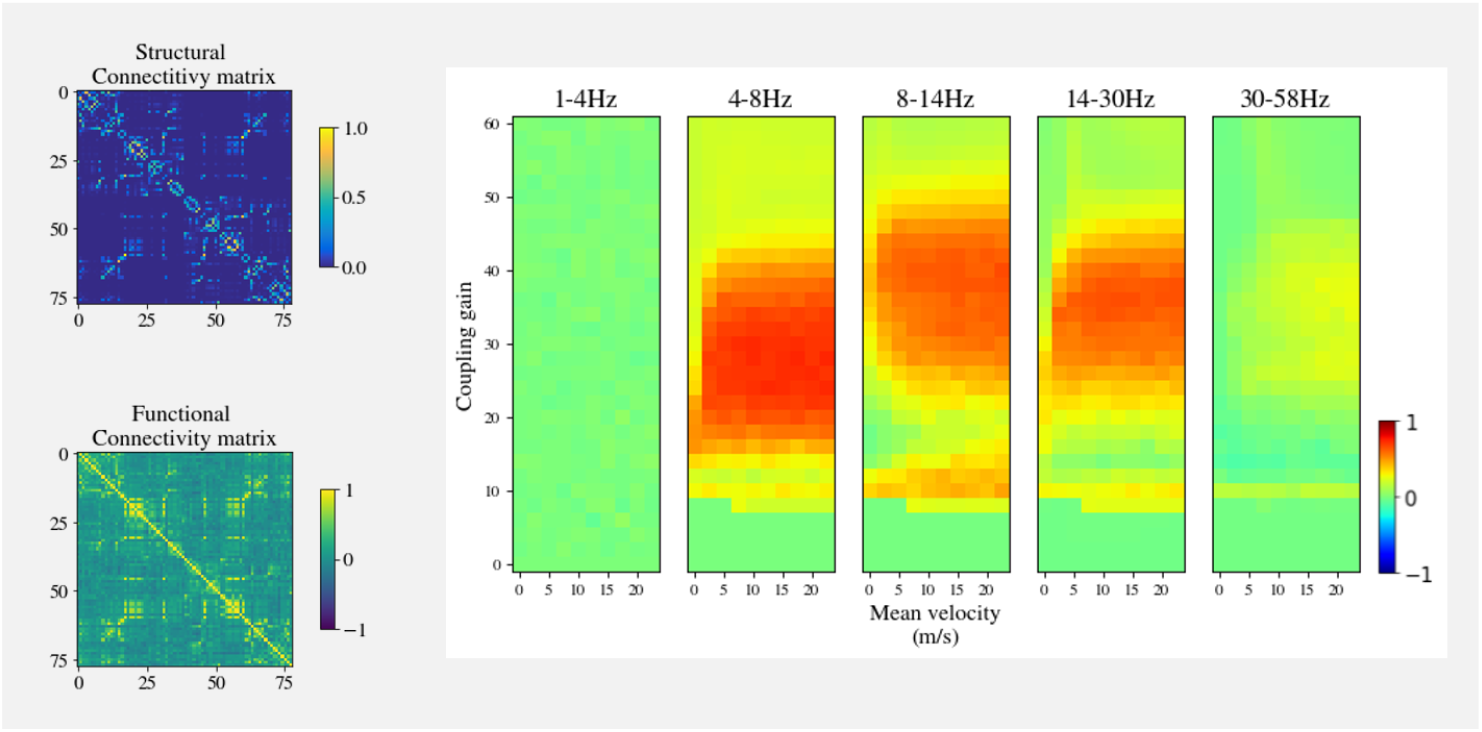
Correlation between structural and HBM functional connectivity. Left: generic HBM structural connectivity matrix and its maximum correlated FC matrix for a pair of model parameters: *G* = 10, *v* = 25 *m/s* in the 4-8 Hz frequency band. Right: Correlation maps between the underlying structural connectivity matrix and the FC matrices extracted from all combinations of model parameters, for different frequency bands.

Moreover, we can observe that the correlation between structural and functional connectivity is more sensitive to the coupling gain *G* than to the mean velocity *v*. The reason for this is that the coupling gain is a bifurcation parameter: as it increases from *G* = 0 to ~ 10, we observe lower background activity. Around *G* = 11, a global bifurcation begins and the system operates in a spiking regime. From *G* = 12 to 55 the model enters a Hopf bifurcation, simulating alpha waves, and, finally, from *G* = 55 onward the NMM returns to a higher background activity (similar to the initial one). For the interested reader, we provide in Appendix C a study of the model’s behavior as *G* is varied. Grimbert and Faugeras (2006), Ableidinger et al. (2017) and Ahmadizadeh et al. (2018) analyze extensively the stability and bifurcations of the Jansen and Rit system and its dependence on different model parameters, including the coupling gain between column connections.

### 3.2. Correlation of HBM and subject-specific FC data

The FC profiles obtained from simulations with different parameters (*G*, *v*) were compared with the FC profiles obtained from real EEG data in different bands for each of the available subjects. In Figure 6a we provide the correlation maps between the HBM derived FC with the FC extracted from the source estimate of Subject 60 in the EC condition in each frequency range. For comparison, Figure 6b) displays the results using a randomized underlying structural connectivity matrix in the generic HBM to further study the effects of the underlying structural connectivity. The distance matrix was also randomized.

**Figure 6:**
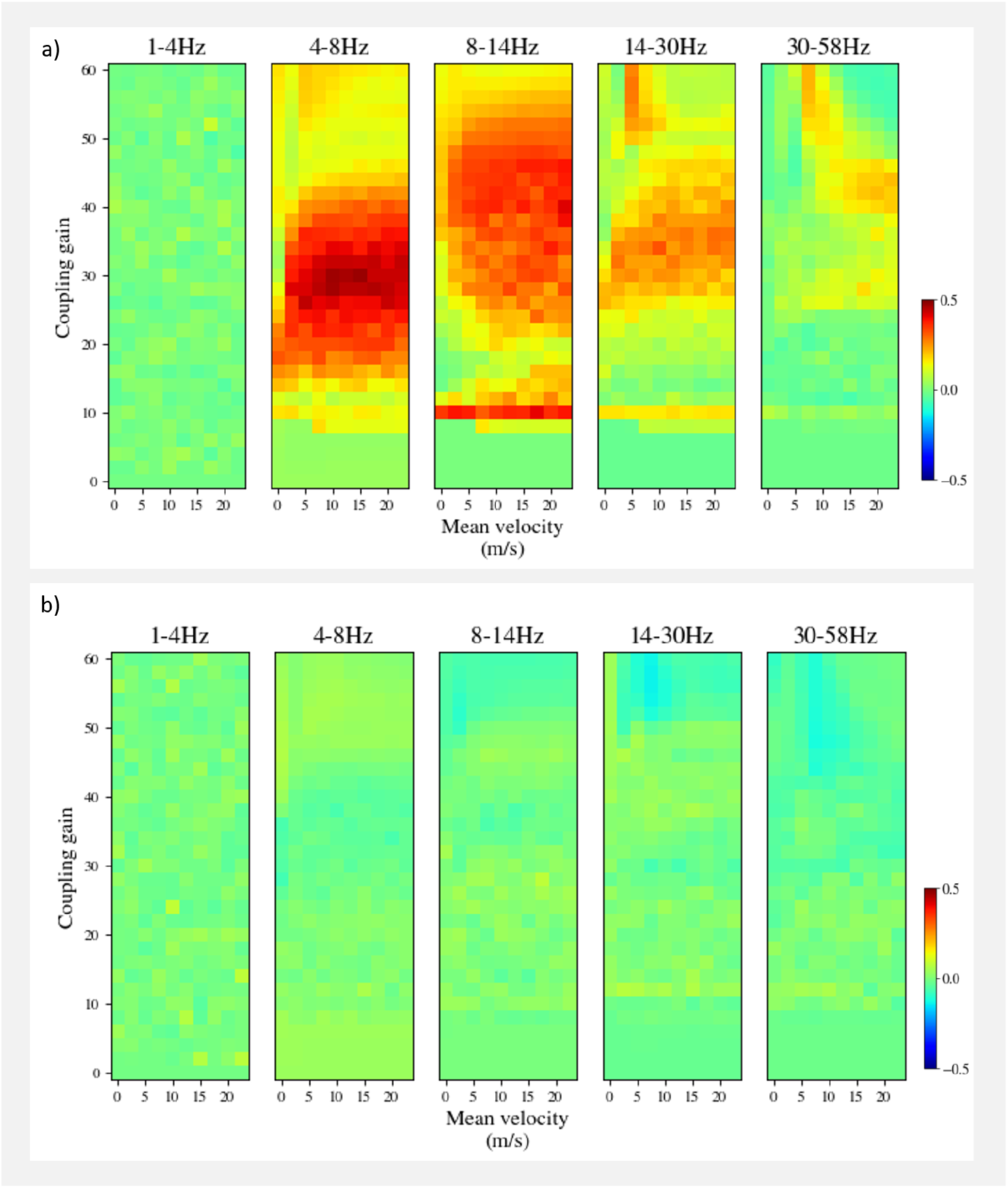
Correlation maps between HBM FC and the subject-specific data-derived FC. **a)** Correlation maps between the simulated FC with the standard underlying connectivity and the experimental ones from Subject 60 (EC condition). **b)** Same as a) with randomized structural and distance connectivity matrices.

In the case without randomization, the model displays the best agreement with real FC data for a similar range of parameters as the ones mentioned in the previous section — where the system operates at the edge of bifurcations. Figure 7 shows the best model parameter agreement, with a Pearson correlation coefficient of *PCC* = 0.5. In the randomized case, there is no significant correlation of HBM FC with the FC profiles extracted from real data (same subject) for any parameter combination. These results highlight the critical impact of the structural connectivity on the outcome. Only the results with a correlation higher than 0.45 have a *p-value<* 0.001, which provides an indication of statistical significance.

**Figure 7:**
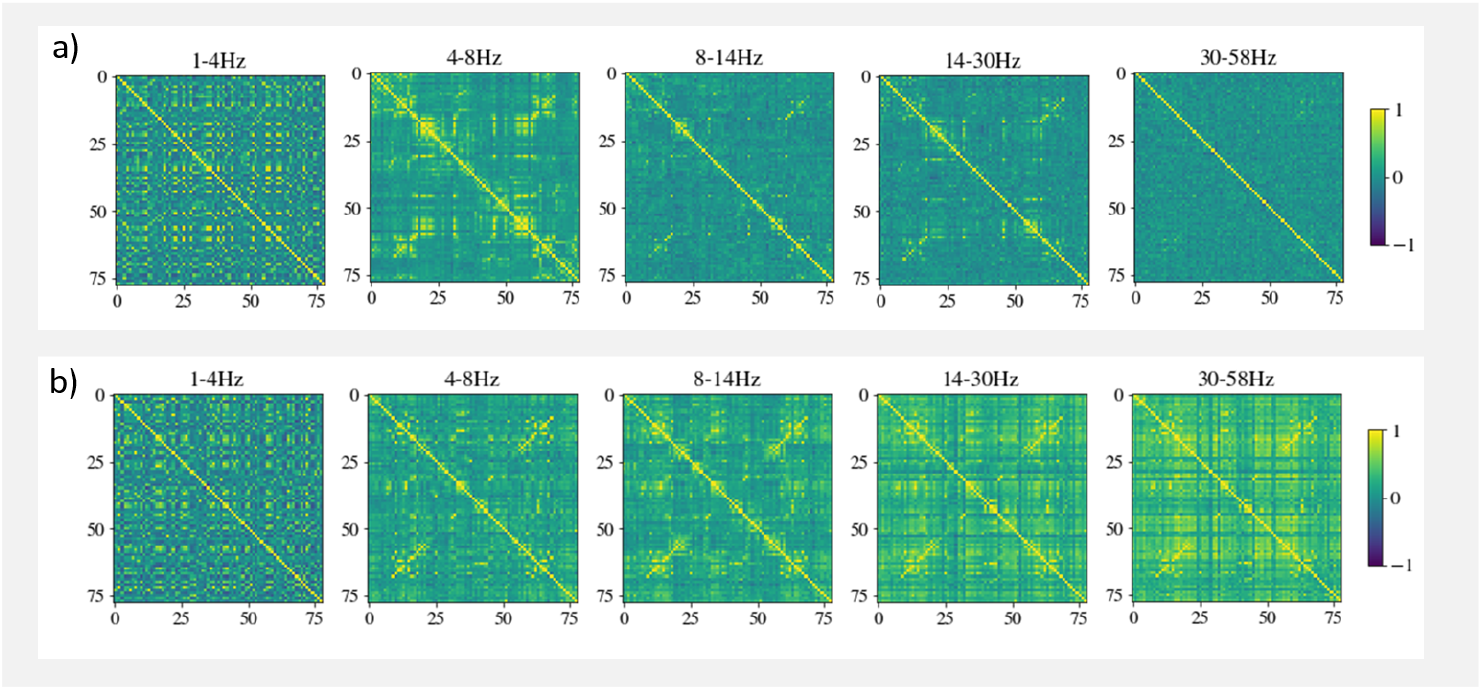
Real and simulated FC profiles, best parameter fit. **a)** simulated FC profiles for the best parameter fit with **b)** real FC profiles: *G* = 30, *v* = 12 m/s, in the frequency range of 4-8 Hz, with a correlation of *PCC* = 0.5

As mentioned in Keilholz et al. (2017), whole brain computational models have shown that the structural connectivity of the brain is a major determinant of the patterns of functional connectivity. However, most of the models perform poorly when confronted with the network dynamics extracted from rs-fMRI (Messé et al., 2014), suggesting that incorporating information obtained with dynamic analysis into the modeling process can serve as a constraint on the types of models and parameters that are appropriate. In consequence, the approach that we have followed in this study could be improved by taking into account the *dyanmic* FC (FCD) matrix, instead of the static one, as explained in Hansen et al. (2015).

### 3.3. Subject and condition variability

We next compare the fit quality for each of the subjects and conditions (eyes open, EO, and eyes closed, EC) to study inter- and intra-subject variability. Figure 8 provides the mean, maximum and minimum correlation between the HBM and each of the subjects for each frequency bands. There are not many significant differences between conditions (EO, EC). This could be due to the choice of the EEG-preprocessing pipeline and its unknown impact on model parameter fit. It remains to be explored if the model can capture other intra-subject conditions (e.g. age, or task performance). Moreover, it can be observed that subject 60 generally correlates more positively with the HBM than subject 55: subject 60 has a significantly higher correlation for delta frequency band (4-8 Hz) and subject 55 has significantly lower correlations for delta (4-8 Hz) and beta bands (14-30 Hz). Further studies should be carried out with a significant number of subjects to test if there is any specific parameter combination capable of clustering a group of individuals (male-female, healthy controls-patients, etc.).

**Figure 8:**
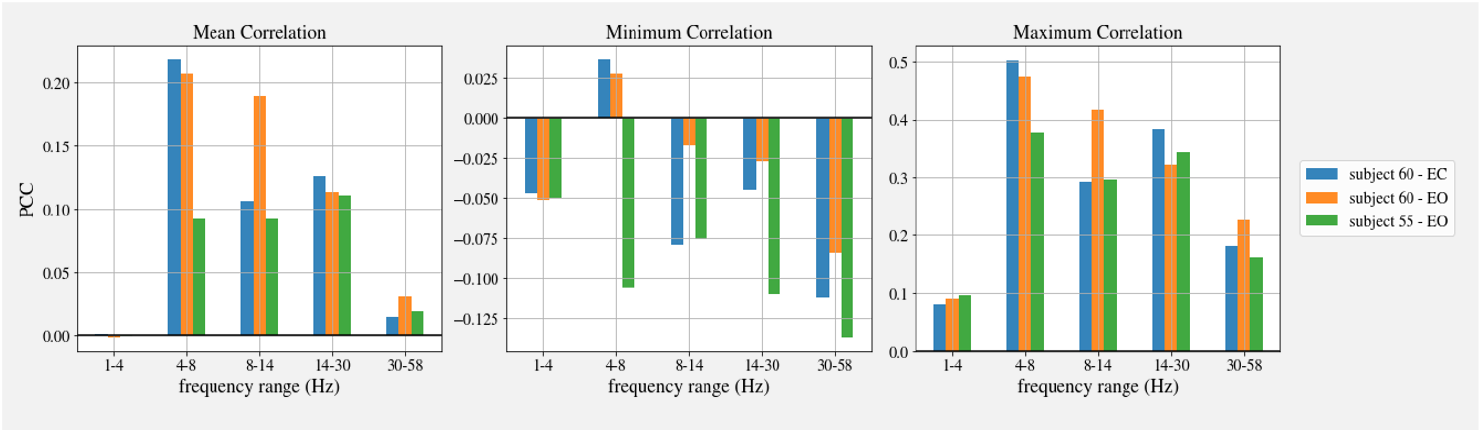
Inter- and intra-subject variability quality of fit study. Mean, minimum and maximum of the cross-correlation coefficient of the EEG-derived FC vs. HBM FC profiles for each of the subjects and frequencies.

## 4. Conclusions

The FC matrices extracted from the HBM proposed in this study display a tight relationship with underlying structural connectivity (Figure 5, *PCC_max_* = 0.74), even higher than shown in previous studies (Cabral et al., 2012). The highest correlations occur when the whole-brain model operates at the Hopf bifurcations of the Jansen and Rit NMM. The increased correlation in this model is also found when comparing real and simulated FC profiles (Figure 6). This seems to be a common feature of these types of models as other studies have observed similar outcomes (Cabral et al., 2011; Deco et al., 2009).

The strong role played by the structural connectivity observed in the mentioned results, together with the fact that model stability is really sensitive to the coupling gain *G*, suggest that the personalization of the model could be significantly improved using subject specific structural connectivity data (DTI).

HBM stability displays lower sensitivity to the mean velocity of transmission for the range of the study (1-20 m/s). These results could be enhanced if instead of relying on the Euclidean distance between the center of masses of each of the AAL regions we used the distance along tracts, which would be physiologically more appropriate. This can be derived from DTI data using tractography algorithms.

A more precise estimation of mean velocity from HBM personalization could provide interesting biomarkers, for example, in multiple sclerosis patients.

### 4.1. Limitations

Several limitations in this study must be mentioned, as they suggest the direction for future improvements.

First, as mentioned in the previous section, results show that the HBM underlying structural connectivity matrix has a clear impact on the functional outcome and the stability of the system is really sensitive to the coupling gain, *G*. Thus, a clear limitation of this study is the lack of individualized DTI data.

We also had to reduce the number of parcellated areas in the cortex because automatic segmentation did not recognize some of them in our subject head models. This fact limits our study since it has been shown that when modeling large scale brain networks, an atlas with at least 140 brain regions produces a good agreement with experimental data (Proix et al., 2016), and we were restricted to use 78 cortical areas. A solution to this issue could be to divide each of the areas into smaller and more homogeneous regions.

Another limitation in our study is associated to the number of recording electrodes in the EEG montage. With 60 EEG electrodes, for example, it is not possible to generate 78 independent areas from cortical mapping. For this reason we had to remove the common mode of the signals instead of orthogonalizing, as it is usually done (Hipp et al., 2012). Moreover, detailed studies such the one in Hassan et al. (2014), which analyze inversion methods and montages, demonstrate a clear improvement in accuracy using weighted minimal norm estimates (similar to the one we are using, considering our restriction on the location of dipoles to a surface) showing the best result when using 180 electrodes.

Further research also needs to be carried out on the effects on the FC profile estimation of the EEG pre-processing pipeline used here. It is unclear, for example, how much impact the interpolation of the bad channels can add to the process.

### 4.2. Outlook

The present study is a proof of concept suggesting that the proposed modeling approach using HBMs has the potential to represent meaningful features of EEG data. It also shows that the pipeline used to extract the functional connectivity and further correlation between simulated and real data is sensitive to inter-subject variability and conditions.

The concept of HBM we propose is, however, more general than what we presented. For example, HBMs could represent biophysical and physiological features using more refined NMMs including other types of neurons as in Merlet et al. (2013) and Wendling et al. (2016). Indeed, the neural mass model (Jansen et al., 1993) used in this study does not fully account for the diversity of neuronal processes occurring at local population level and in the coupling between distant populations. In particular, it only features one sub-population of GABAergic interneurons (somatostatin-positive (SST) neurons) mediating slow GABAa inhibitory PSPs onto pyramidal cells. Recent findings indicate that two other types of GABAergic interneurons are involved both in the activity locally-generated by each brain region and in the communication between distant regions: parvalbumin-positive (PV) and Vasoactive Intestinal Peptide (VIP) interneurons. PV “fast-spiking” GABAergic interneurons play a key role in the generation of local gamma activity (Pyramid-Interneuron Gamma sometimes referred to as the “ping” circuit) (see review Hu et al. (2014)). In contrast, VIP interneurons which can be activated from distant pyramidal cells are determinant for information transfer and processing in large scale networks (see review Cardin (2018)). These results suggest that, in addition to collateral excitation among pyramidal cells in the cortex, feedforward and feedback inhibition also play a role in complex network integration and segregation operations. Future NMM developments can likely account for the specificity of these circuits and thus better reproduce the dynamics of local and global brain activity, the counterpart being an increased number of connectivity parameters and thus a more difficult model parameter identification problem.

Although in the present study a template HBM was personalized from the integration of averaged structural connectivity data (DTI), individual anatomical data (MRI) and EEG, we can envision the assimilation of other types of data. Event related potentials (as in Molaee-Ardekani et al. (2013)), magnetoencephalograpy (MEG), stereo-EEG (depth-EEG recorded with depth electrodes implanted under stereotaxic conditions, SEEG) or transcranial magnetic stimulation combined with EEG (TMS-EEG), for example, can provide valuable information on cortico-cortical connectivity. Fitting such data in a template HBM could widen the scope and precision of parameters to be estimated.

Since they can readily represent biophysics and physiology in a unified manner, HBMs can provide insights into the effects of brain stimulation techniques mediated by macro or mesoscale electric fields, such as tCS or TMS (Ruffini et al., 2013; Merlet et al., 2013; Molaee-Ardekani et al., 2013; Muldoon et al., 2016; Kunze et al., 2016), or drugs (Kurbatova et al., 2016; Liang et al., 2015). This can be achieved as the individual level, and therefore could lead to the development of optimization and personalized treatment strategies. Similarly, HBMs offer a tool to study the role of *ephaptic* interactions in neurodynamics, i.e., non-synaptic neighboring interactions mediated by endogenous electric fields which occur at relatively small scales and which may provide a clue for explaining neurophysiological network effects in the human brain (Jefferys, 1995; Anastassiou et al., 2011). This is of particular interest, not the least because such a mechanism represents a feedback loop between electromagnetism and neurodynamics in the human brain that can be studied at mesoscales.

However, given that the clinically relevant aspects of tCS or transcranial magnetic stimulation (TMS) revolve around their long term effects, HBMs need to represent another crucial physiological aspect, namely neuroplasticity. The integration of plastic effects in HBMs entails the proper representation of rules for the dynamics of connectivity parameters. This will be key step for the use of HMBs for analysis and optimization of neuro-modulatory techniques that rely on such plastic effects for their therapeutic value. In order to do so, the relevant data will need to be collected. For example, tCS can be combined with TMS to study brain plasticity (Nitsche and Paulus, 2000) through the analysis of motor evoked potentials (MEPs). This phenomenon, or others such as those implicated in repetitive transcranial magnetic stimulation (rTMS), can also be represented by HBMs and thus provide the means to fix parameters related to plasticity.

Finally, since neural mass models can simulate physio-pathological neuronal signals such as those in epilepsy (Wendling et al., 2016; Wendling and Chauvel, 2008; Cabral et al., 2012), HBMs built on them can be personalized to represent phenomena in the human brain such as epileptic spikes and epileptogenic networks (Bartolomei et al., 2017), or more generally, the combination of alterations at the microscale (neuronal level, such as excitation/inhibition balance) and at larger scales (connectivity), which are thought to underlie neuropsychiatric disorders. The availability of such virtual brain models will provide a powerful basis for the development of optimized, personalized treatment and more effective neuromodulatory interventions, representing a significant step forward towards mechanistically-grounded, quantitative therapeutics in neuropsychiatry.

## Acknowledgements

This research was supported in part by the Future Emerging Technologies Open Luminous project (H2020-FETOPEN-2014-2015-RIA under agreement No. 686764) as part of the European Union’s Horizon 2020 research and training program 2014–2018.

## Appendix A. Jansen and Rit NMM

Jansen and Rit (Jansen and Rit, 1995) developed a NMM of cortical columns based on the thalamus model of Lopes da Silva et al. (1974)and on the post-synaptic-potential (PSP) transformation function that van Rotterdam et al. (1982) designed. This model simulates a population of pyramidal cells located in layer V of the cortex that receive inhibitory inputs from a sub-population of interneurons, dendrite targeting cells. They also receive excitatory feedback either from a population of excitatory interneurons or other pyramidal cells located in the same cortical column.

Each of the populations is characterized by two state variables: the average membrane potential, *y*(*t*), and average firing rate, *x*(*t*). They are also characterized by two different transformation functions that convert one state variable into the other, the pre-synaptic and post-synaptic functions. These functions reproduce the two essential characteristics of real post-synaptic potentials: shape and excitatory/inhibitory ratios. All the function parameters, their physiological meaning and the default values are summarized in Table A.1.

**Figure A.1:**
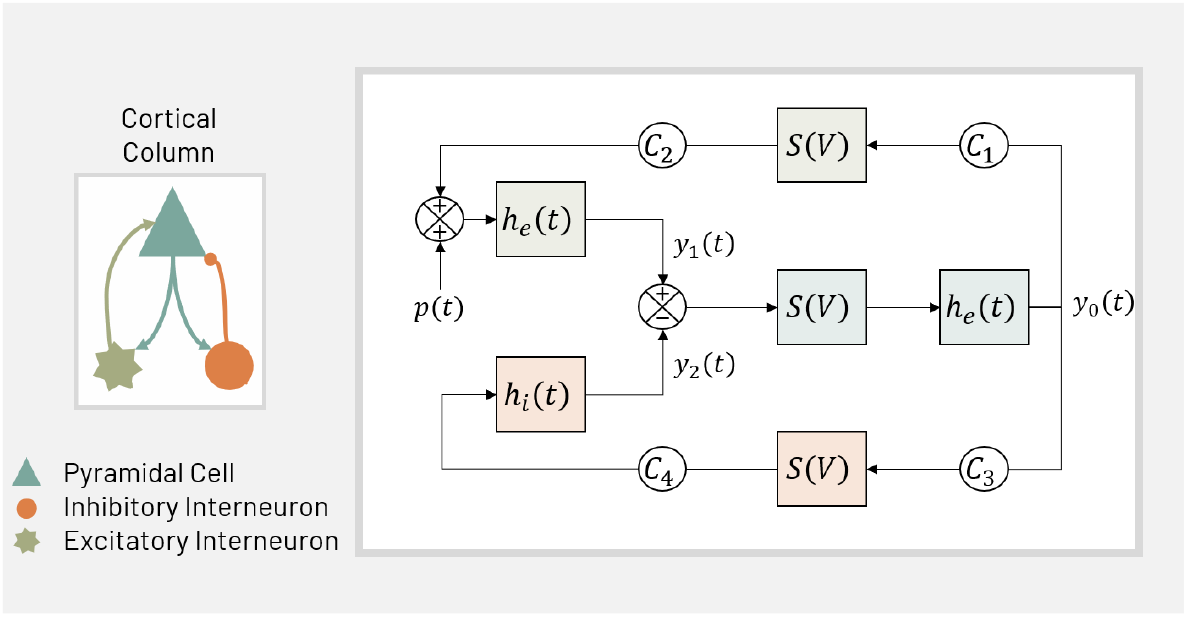
Diagram of Jansen and Rit model for a cortical column. Each color and shape represent one type of column population: pyramidal (blue), excitatory interneuron (green) and inhibitory interneuron (orange).

**Table A.1:** Jansen and Rit neural mass model parameters, with their associated physiological interpretation and default values.

| Parameter | Description | Value |
| --- | --- | --- |
| $A$ | Average excitatory synaptic gain | 3.25 mV |
| $B$ | Average inhibitory synaptic gain | 22 mV |
| $a$ | Time constant of excitatory PSP | 100 s<sup>-1</sup> |
| $b$ | Time constant of inhibitory PSP | 50 s<sup>-1</sup> |
| $C_n$ | Average number of synaptic contacts | $C_1 = 135$<br>$C_2 = 0.8 \cdot C_1$<br>$C_3 = 0.25 \cdot C_1$<br>$C_4 = 0.25 \cdot C_1$ |
| $v_0$ | Potential when half of the maximum firing rate is achieved | 6 mV |
| $e_0$ | Half of the maximum firing rate | 2.5 s<sup>-1</sup> |
| $r$ | Slope of the pre-synaptic function at $v_0$ | 0.5 mV<sup>-1</sup> |
| $p(t)$ | Excitatory input noise (mean Gaussian white noise) | $\mu = 90$ s<sup>-1</sup><br>$\sigma = 30$ s<sup>-1</sup> |

The pre-synaptic function, *σ*(*v*), introduces a nonlinear component that transforms the average membrane potential of a population (in mV) into average firing rate:

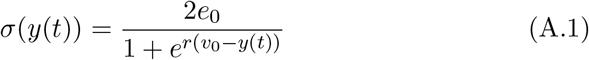

The post-synaptic function converts the average firing rates into an average PSP. This filtering process is introduced as a second order differential linear transformation:

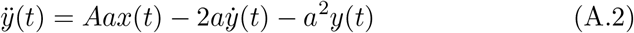

where *x*(*t*) is the output of the sigmoid function (average firing rate). Equivalently, a system of two equations can be written:

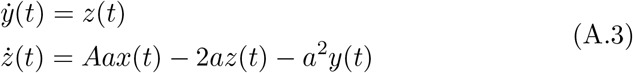

The Jansen and Rit NMM of cortical columns has three main state variables, the average membrane potential of each of the subpopulations, *y*_0_ for the pyramidal cells and *y*_1_, *y*_2_ for the excitatory and inhibitory interneurons, respectively. The main output of the model is the average membrane potential of the pyramidal cell population, as the sum of the potential of these cells is thought to be the source of the potential recorded in the EEG. Then, the equations describing the Jansen and Rit model are the following:

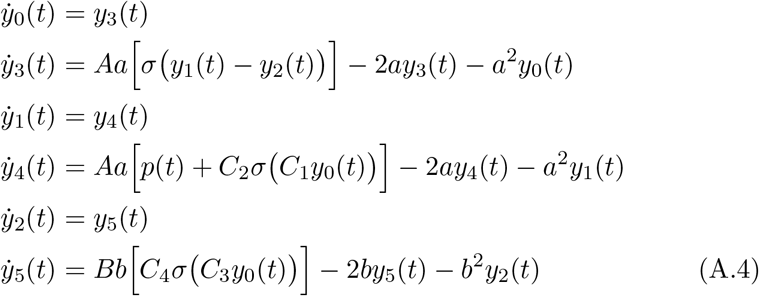

## Appendix B. Cortical Regions of the AAL atlas

In Table B.1 there is a list of the AAL(Tzourio-Mazoyer et al., 2002) cortical areas extracted from Achard (2006). The order of the cortical regions used in this paper in structural or functional connectivity matrices is represented in Figure B.1.

**Figure B.1:**
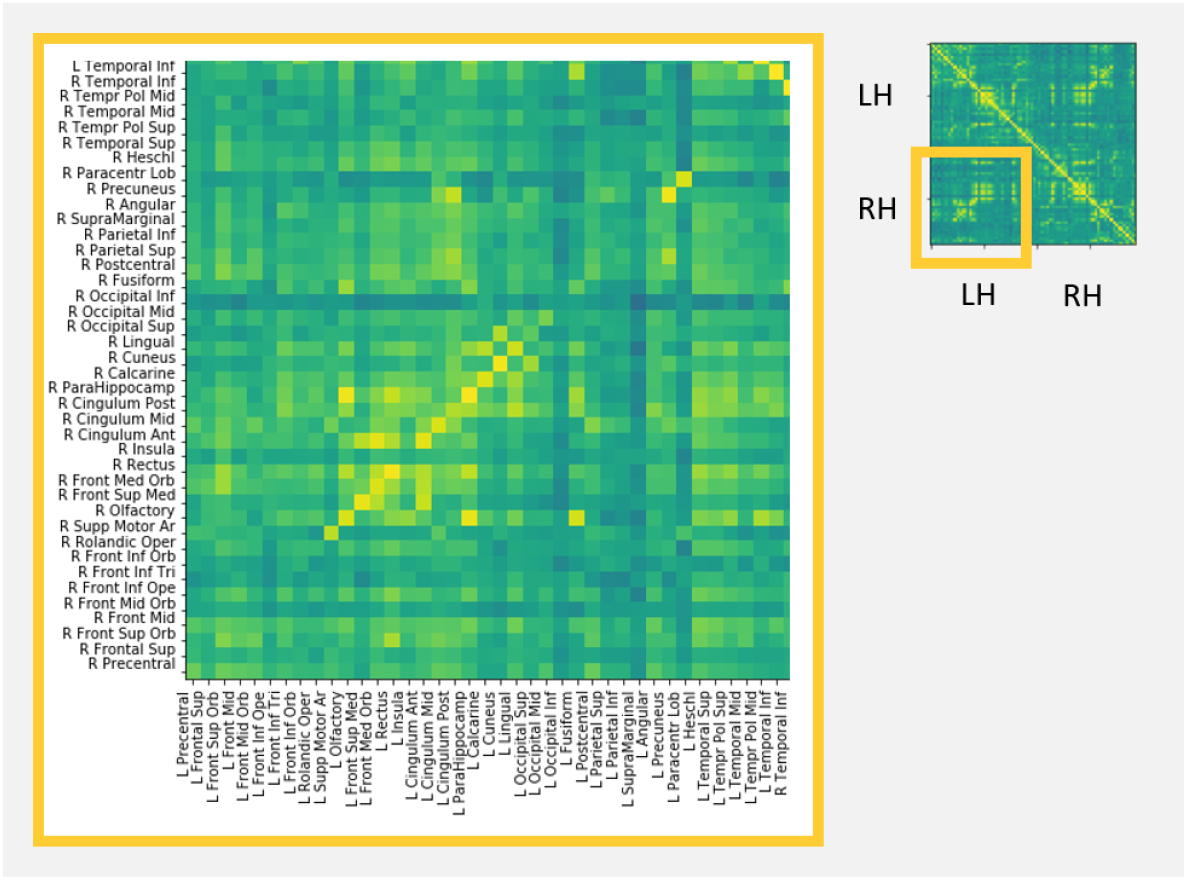
Ordering of the brain areas in a structural or functional connectivity matrix.

**Table B.1:** Cortical Areas in the AAL template of one hemisphere (Tzourio-Mazoyer et al., 2002; Achard, 2006; Gong et al., 2009)

| # | Area label | # | Area label |
| --- | --- | --- | --- |
| 1 | Precentral | 21 | Cuneus |
| 2 | Frontal Sup | 22 | Lingual |
| 3 | Frontal Sup Orb | 23 | Occipital Sup |
| 4 | Frontal Mid | 24 | Occipital Mid |
| 5 | Frontal Mid Orb | 25 | Occipital Inf |
| 6 | Front Inf Ope | 26 | Fusiform |
| 7 | Front Inf Tri | 27 | Postcentral |
| 8 | Front Inf Orb | 28 | Parietal Sup |
| 9 | Front Rolandic Oper | 29 | Parietal Inf |
| 10 | Supp Motor Ar | 30 | SupraMarginal |
| 11 | Olfactory | 31 | Angular |
| 12 | Front Sup Med | 32 | Precunneus |
| 13 | Front Med Orb | 33 | Paracentr Lob |
| 14 | Rectus | 34 | Heschl |
| 15 | Insula | 35 | Temporal Sup |
| 16 | Cingulum Ant | 36 | Temporal Pol Sup |
| 17 | Cingulum Mid | 37 | Temporal Mid |
| 18 | Cingulum Post | 38 | Temporal Pol Mid |
| 19 | ParaHippocamp | 39 | Temporal Inf |
| 20 | Calcarine |  |  |

## Appendix C. Appendix: Study of bifurcations for the coupling gain

When the input noise *p*(*t*) is varied smoothly the eigenvalues of the fixed points of the system of equations that describe the Jansen and Rit NMM (Jansen et al., 1993) move across the complex plane leading to global and Hopf bifurcations (see Figure C.1b, bifurcation diagram taken from Grimbert and Faugeras (2006),a deep study on the bifurcations that characterize the Jansen and Rit NMM). To summarize it, Hopf bifurcations generate the alpha activity seen in Jansen et al. (1993) (Figure C.1, blue lines) and the global bifurcations generate the spike-like epileptic activity that Wendling and colleges Wendling et al. (2016); Wendling and Chauvel (2008)use to simulate the SEEG recordings in epileptic patients (Figure C.1, yellow lines). When connecting multiple columns (e.g. generating the HBM), we have observed that our system stability also changes in a similar way than when varying the input noise *p*. See David and Friston (2003); Kunze et al. (2016); Ableidinger et al. (2017); Ahmadizadeh et al. (2018)for an extended study on the effect of the coupling factor in different networks of Jansen and Rit models of cortical columns. In Figure C.1a we can see the effect of increasing values of the coupling gain *G* on some random nodes of our HBM.

**Figure C.1:**
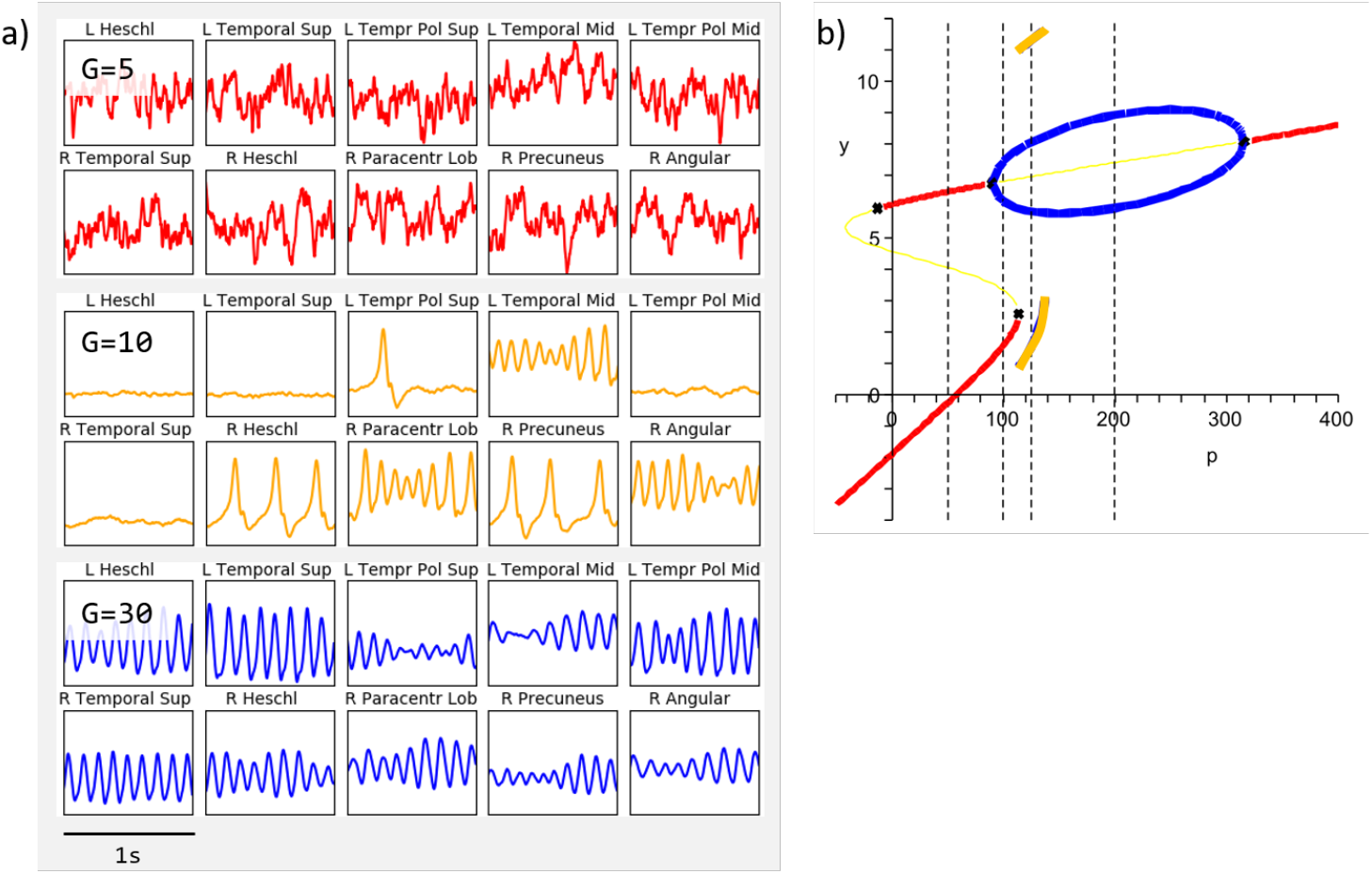
a) Study of the behaviour of the HBM for increasing values of the coupling gain *G* = 5 (red), *G* = 10 (yellow), and *G* = 30 (blue). b) Bifurcation diagram of a Jansen and Rit NMM for the bifurcation parameter *p*, adapted from Grimbert and Faugeras (2006) (Figure 24, p. 29) to reflect corresponding colors in a).

